# The accessory protein CvnF8 modulates histidine kinase activity in an Actinobacterial G protein system in *Streptomyces coelicolor*

**DOI:** 10.1101/2025.07.03.663114

**Authors:** Luis M. Cantu Morin, Kilian Dekoninck, Kyung-Yoon Min, Matthew F. Traxler

## Abstract

Conservons are regulatory systems found in bacteria of the phylum Actinomycetota. These regulatory systems are composed of four core proteins: a sensor histidine kinase-like protein (CvnA homolog), an MglA-type roadblock protein (CvnB homolog), a protein containing a domain of unknown function (CvnC homolog), and small Ras-like GTPase (CvnD homolog). Based on their conserved small GTPase components and their phylogenetic distribution, we propose that conservons should be known as <u>A</u>ctinobacterial <u>G p</u>rotein <u>s</u>ystems (AGPSs). The signal transduction path through AGPSs remains poorly understood, and some AGPSs have additional accessory proteins (CvnE and CvnF homologs) of unknown function. In this work, we show that AGPS accessory proteins are present when the cognate histidine kinase protein (CvnA homolog) lacks an extracytoplasmic sensory domain. It was previously shown that the Cvn8 AGPS of *Streptomyces coelicolor* controls expression of multiple pathways for specialized metabolism and that the Cvn8 AGPS contains an accessory protein, CvnF8. Through protein modeling, we found that CvnF8 may share an interaction interface with the histidine kinase CvnA8, prompting the hypothesis that CvnF8 may serve as a modulator of CvnA8 activity. We found that in a purified system, CvnF8 strongly stimulated the ATPase activity and autophosphorylation of CvnA8. Taken together, these findings indicate that CvnF family accessory proteins likely serve as sensors and/or modulators of histidine kinases of AGPSs found broadly in Actinomycetota.

**Importance Statement:** Many lineages of bacteria in the phylum Actinomycetota contain conserved operons (conservons) that encode an unusual type of regulatory system whose function is poorly understood. These lineages include pathogens such as *Mycobacterium tuberculosis* and members of the genus *Streptomyces* that produce valuable natural products. These regulatory systems are composed of four proteins, including a sensor histidine kinase, a small Ras-like GTPase, a likely GTPase activating protein, and a protein containing a domain of unknown function. Given this composition, we propose that these regulatory modules be known as Actinobacterial G protein systems (AGPSs). We show that some AGPSs include accessory proteins that are only found with partner histidine kinases that lack sensory domains. We demonstrate that one such accessory protein can control the activity of its cognate histidine kinase. Together this work indicates that these CvnF-family accessory proteins likely serve as sensory inputs for AGPSs found broadly in Actinomycetota.

## Introduction

Conservons are four-gene operons that were initially discovered in the genome of the model actinomycete *Streptomyces coelicolor*, in which they appear thirteen times (1). Genomic data has since shown that conservons are found broadly across organisms in the phylum Actinomycetota, including pathogens such as *Mycobacterium tuberculosis* (2, 3). Subsequent experimental work has suggested that conservons encode signal transduction systems that share features of both bacterial two-component signaling systems and eukaryotic-style GTPase signaling cascades (4–7). Together, these observations indicate that conservons are an important class of signal transduction systems in actinobacteria whose human relevance ranges from pathogens to producers of important natural products.

In *Streptomyces coelicolor*, the four syntenic core genes of conservons are conventionally labeled *cvnA*-*cvnD, 1-13* (1). These genes encode a predicted sensor histidine kinase (CvnA homolog), an MglB-type roadblock protein (CvnB homolog), a small protein with a domain of unknown function (CvnC homolog), and a Ras-like GTPase (CvnD homolog) (1, 2, 4). Given the putative functions of these four conserved proteins, and the growing evidence that they function together in signaling cascades, we propose that conservons should be understood as Actinobacterial G protein systems (AGPSs). In addition to the four core components, a subset of AGPSs also have a fifth accessory gene in the operon which typically fall into one of two categories, encoding either a cytochrome P450 (*cvnE*-type homolog) (1) or a GAF domain-containing protein (*cvnF*-type homolog) (6). Importantly, neither of these accessory protein types has been assessed at the biochemical level.

While our current understanding of how AGPSs function and the processes they regulate is fragmentary, a picture of signal flow through these systems is beginning to emerge. A previous study found that the *S. coelicolor* Cvn8 system is a global regulator of genes involved in specialized metabolism during interspecies interactions, with *cvn* mutants showing altered gene expression in pathways for the biosynthesis of actinorhodin, prodiginines, and coelimycins (6).

In the same report, it was shown that the Cvn8 system normally represses transcription of an adjacent lanthipeptide biosynthetic gene cluster whose theoretical peptide product was recently termed lanthiphage_Sco (8). In addition to the AGPS core genes *cvnA8-D8*, the *cvn8* operon includes *cvnF8*, which encodes a GAF domain-containing accessory protein.

Transcriptomic analysis of mutants lacking individual genes of the Cvn8 system suggested that this system could be divided into upstream and downstream components (6). Specifically, it was hypothesized that CvnA8 and CvnF8 function in the upper, sensory input section of the pathway, while CvnC8 and CvnD8 are involved in downstream signaling. It was also hypothesized that CvnB8 could serve as a key connection between CvnA8/F8 and CvnD8/C8.

Work on another AGPS, the Mfp system, which is involved in fluoroquinolone resistance in *Mycobacteria*, demonstrated that the CvnB homolog MfpD serves as a <u>G</u>TPase <u>a</u>ctivating <u>p</u>rotein (GAP) for its cognate GTPase, MfpB (5). Further work showed that the CvnC homolog MfpC likely functions as a guanine nucleotide exchange factor (GEF) for MfpB (7). Taken together, these studies point to a model wherein Actinobacterial G protein systems contain a sensory protein (CvnA homolog) that indirectly passes a signal to a small GTPase (CvnD homolog), likely by modulating the activities of a GAP (CvnB homolog) and/or GEF (CvnC homolog). Thus, while some functionality of the CvnB/C/D components of Actinobacterial G protein systems is starting to come into focus, the sensory input components of these pathways, i.e. how the histidine kinases and their possible accessory proteins function, remains largely unexplored.

In this work we sought to understand the role of the CvnF8 accessory protein in the Cvn8 AGPS. As a starting point, we mapped the occurrence of Actinobacterial G protein systems and accessory proteins across 25 orders within the phylum Actinomycetota. This analysis revealed a key evolutionary pattern; the accessory proteins CvnF and CvnE only accompany CvnA homologs (histidine kinases) that lack an extracellular sensory domain. We show that CvnF8 interacts directly with CvnA8, stimulating its ATPase activity and autophosphorylation *in vitro*. Finally, we show through epistasis analysis that CvnF8 and CvnA8 exhibit functional dependence within the Cvn8 AGPS cascade. In sum, these results demonstrate that the CvnF class of GAF domain-containing accessory proteins are modulators of CvnA-type histidine kinases found in a wide range of Actinobacterial G protein systems.

## Results

### Actinobacterial G protein Systems (AGPSs) are found widely across the Actinomycetota

Actinobacterial G protein systems (AGPSs) are found in Actinomycetota, but their phylogenetic distribution within this phylum remains only partially resolved. To investigate whether AGPSs are universally present or have undergone lineage-specific divergence, we constructed a species tree using a concatenated alignment of fifteen ribosomal proteins from 485 genomes selected from the ActDES database, a curated, non-redundant collection of genomes from Actinomycetota (9). Orthologous gene families corresponding to AGPS components were identified using OrthoFinder, and their presence or absence across genomes was mapped onto the species tree as colored heatmap tracks (Fig. 1). In this analysis, a genome is considered positive for core AGPS genes (*cvnA-D*) if a subset of these genes occurs in synteny –defined as being at a specific gene-distance from a *cvnA* homolog: *cvnB* one gene away from *cvnA*, cvnC two genes away from *cvnA*, *cvnD* three genes away from *cvnA*, and *cvnE* or *cvnF* four genes away from *cvnA*. We also counted the number of AGPSs present in each genome, shown as a grey bar at the right side of the figure.

**Fig. 1.**
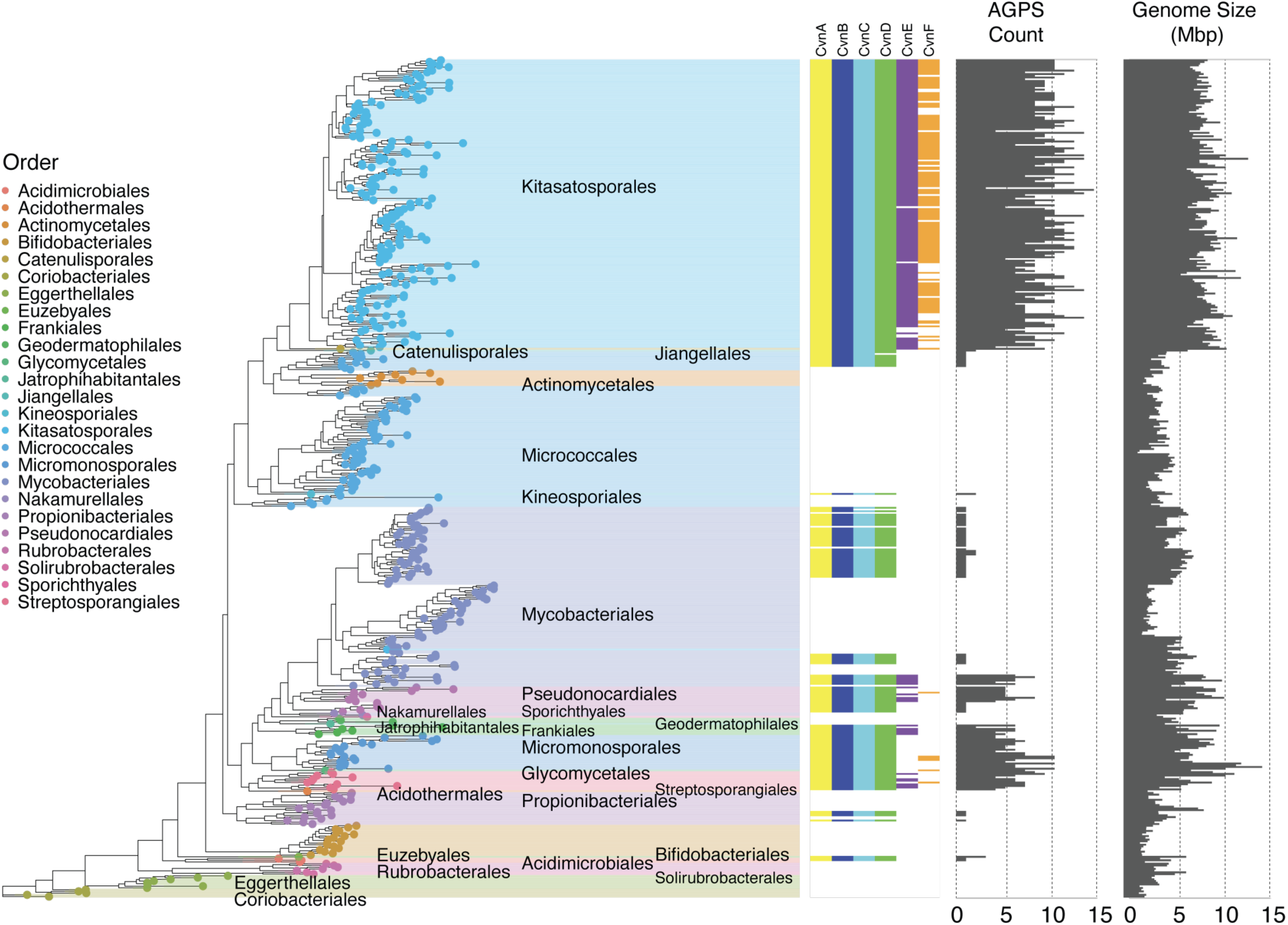
Actinobacterial G protein systems (AGPSs) are distributed across Actinomycetota. A species tree of 485 Actinomycetota genomes, highlighting the distribution of AGPSs across the phylum. AGPSs are detected in 289 genomes. The heatmap tracks indicate the presence of orthogroups CvnA, CvnB, CvnC, and CvnD, found in synteny. Orders within Actinomycetota are labeled along the tree. Two additional tracks summarize genome features: the AGPS Count bar plot indicates the total number of AGPS operons per genome, and the Genome Size bar plot reports the total genome size (in megabase pairs, Mbp).

This analysis shows that AGPSs are widely distributed across at least fourteen orders within the Actinomycetota. Most orders whose members typically have large genomes (i.e. >7 Mb) contain 5-10 AGPSs, with some species having as many as fourteen. A significant positive correlation (Pearson’s r = 0.68) was observed between AGPS count and genome size (Fig. S2), suggesting that genomes with larger coding capacity tend to harbor more AGPS loci. These orders include the Kitasatosporales (which contains the genus *Streptomyces*), Micromonosporales, Pseudonocardiales, Catenulisporales, Streptosporangiales, and Frankiales. A subset of Mycobacteriales also typically contain at least one AGPS. These include both *M. tuberculosis* and nontuberculous pathogenic *Mycobacteria*. In most cases, the single AGPS represents the Mfp system involved in fluoroquinolone resistance (5). Lastly, we also find that several basal orders such as the Acidimicrobiales, Euzebyales, and Propionibacteriales contain AGPSs, though the number per genome is typically one.

We also sought to examine the distribution of the AGPS accessory proteins CvnE and CvnF. We found that AGPS operons with a *cvnE* or *cvnF* accessory protein orthologs are limited to those clades whose members have large genomes. The limited presence of *cvnE* and *cvnF* orthologs in these orders suggests that these accessory components may have evolved later as lineage-specific adaptations.

### *cvnF* and *cvnE* accessory genes are associated with CvnA proteins that lack sensory domains

Previous studies (1, 4), and protein domain analyses showing that CvnA homologs share key features with histidine kinases, suggest that they serve as the primary sensors of AGPSs. Broadly speaking, these features include transmembrane domains, sensory domains, HAMP domains (involved in intra-protein signal transduction), and ATPase domains.

We sought to systematically examine CvnA domain architecture across this protein family. To do so, we constructed a gene tree of the CvnA orthogroup with 2,175 gene sequences from across the *Actinomycetota* (Fig. 2A). Labeling positions on the tree with the names of twelve CvnA orthologs found in the *S. coelicolor* genome showed that a single organism can harbor a diverse set of these sensory proteins.

**Fig. 2.**
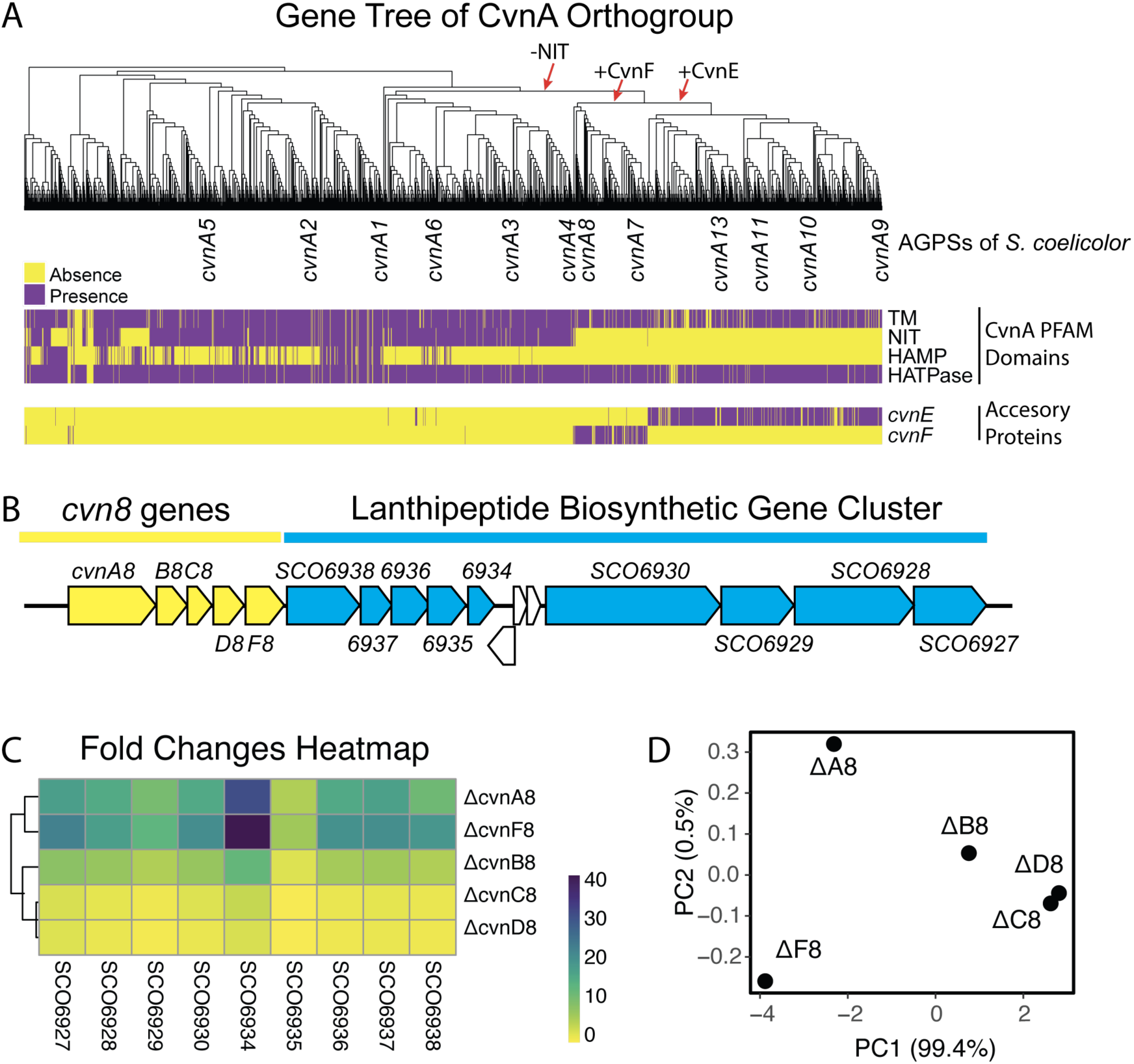
Modularity of CvnA and its association with accessory proteins and lanthipeptide gene expression patterns in Cvn8 mutants. (A) Gene tree of CvnA homologs across Actinomycetota, annotated with PFAM domain architecture (TM, NIT, HAMP, HATPase) and presence of neighboring accessory proteins from the CvnE and CvnF orthogroups. The heatmap reflects domain composition and syntenic co-occurrence of CvnF and CvnE, suggesting functional modularity and potential co-evolution. (B) Schematic of the lanthipeptide biosynthetic gene cluster (*lan* BGC) under the regulatory control of the Cvn8 Actinobacterial G protein system in *Streptomyces coelicolor*. (C) Heatmap showing the average transcript fold-change of *lan* BGC genes measured via NanoString nCounter analysis, highlighting derepression of target genes in *ΔcvnA8* and *ΔcvnF8 mutants*. (D) Principal component analysis (PCA) of the expression data from (C), illustrating transcriptional similarity between *ΔcvnA8* and *ΔcvnF8* strains.

A heatmap track shows the presence or absence of four PFAM domains within CvnA proteins including transmembrane domains (TM), Nitrate Sensing family domains (NIT), HAMP domains, and Histidine Kinase ATPase (HATPase) domains. This analysis shows that CvnA proteins exhibit variable modularity, with some lineages retaining all canonical domains, while others have entirely lost NIT sensory domains. We interpret the structural variability seen across the clades of CvnA proteins as reflecting the variety of inputs that this protein family has adapted to sense.

We hypothesized that modularity in CvnA domain architecture might coincide with the presence of CvnE and CvnF accessory proteins. To investigate this possibility, we checked the presence/absence of syntenic CvnE and CvnF accessory proteins with a separate heatmap track. This analysis revealed a clear correlation: CvnE or CvnF proteins are overwhelmingly found in synteny with CvnA proteins that lack sensory domains. This correlation implies that CvnE/CvnF accessory proteins may function as sensory inputs and/or modulators of activity for a specific set of CvnA orthologs.

### Mutations in *cvnA8* or *cvnF8* lead to similar gene expression patterns

The above analyses suggest that the function of CvnE and CvnF accessory proteins are likely linked to their cognate CvnA proteins. This prompts the prediction that mutant strains lacking an accessory protein, such as a CvnF, might share similar phenotypes with strains lacking the cognate CvnA protein. To investigate this possibility, we focused on the Cvn8 system in *Streptomyces coelicolor*, encoded by genes SCO6943-SCO6940.

A previous study (6) demonstrated that the Cvn8 system normally represses expression of an adjacent lanthipeptide biosynthetic gene cluster (hereafter called the *lan* gene cluster, illustrated in Fig. 2B). The Cvn8 system also contains the accessory protein CvnF8 (SCO6939). To both verify the role of the Cvn8 system in repression of the *lan* gene cluster and check the contribution of CvnA8 and CvnF8 to this repression, we measured expression of nine genes in the *lan* gene cluster in Cvn8 system mutants from 3-day old patches of *S. coelicolor* grown on ISP2 agar using the NanoString nCounter analysis platform.

Deletion of either *cvnA8* or *cvnF8* led to a similar strong derepression (7.0 - 41.8-fold higher expression) of genes in the *lan* biosynthetic cluster (Fig. 2C). Hierarchical clustering (via euclidean distance) of gene expression patterns across all the individual mutants (*cvnA-F8*) showed that the expression patterns in the *ΔcvnA8* and *ΔcvnF8* strains grouped together (see dendrogram at left in Fig. 2C). This similarity is also evident in a PCA analysis, which showed that the expression patterns in the *ΔcvnA8* and *ΔcvnF8* strains were most similar to one another on the PC1 axis, which accounted for 99.4% of the transcriptional differences seen across all the mutant strains (Fig. 2D). Additionally, we note that expression patterns in the *ΔcvnA8* and *ΔcvnF8* strains contrasted the patterns seen in the *ΔcvnC8* and *ΔcvnD8* strains, which showed a lower degree of derepression. The *ΔcvnB8* strain showed an intermediate level of derepression. Taken together, the similar expression patterns in the *ΔcvnA8* and *ΔcvnF8* strains are consistent with a model in which CvnA8 and CvnF8 function together within the Cvn8 signaling cascade.

### Mutual Information Analysis identifies co-evolving residues at a predicted CvnA8/CvnF8 interface

To consider whether CvnF proteins might function as accessory proteins to CvnA histidine kinases, we tested whether residues in CvnA and CvnF proteins have coevolved, which would suggest functional or physical interactions. We hypothesized that CvnA and CvnF proteins must not only interact but also exhibit evolutionary constraints at specific sites, particularly at their protein interfaces. To test this, we calculated mutual information (MI) scores for residues across cognate CvnA and CvnF pairs, identifying coevolving residue pairs.

Mutual information analysis revealed significant coevolution between CvnA and CvnF proteins, with high MI scores (>0.7) indicating strong residue-residue dependencies. When comparing true CvnA-CvnF protein pairs (conjugate pairs) to randomized pairs, we observed an enrichment of coevolving residues in true pairs, supporting a functional relationship between cognate CvnA and CvnF proteins (Fig. 3A). Further analysis of MI scores revealed two distinct categories: within-protein coevolution (residues evolving together within CvnA or within CvnF proteins) and between-protein coevolution (residues evolving together across CvnA and CvnF conjugate pairs), with the latter suggesting a direct or allosteric interaction between the two proteins. This analysis showed that residues within CvnA proteins that have high between-protein coevolution with resides in CvnF proteins (MI > 0.7) are enriched in the HATPase domains of CvnA homologs (Fig. 3B).

**Fig. 3.**
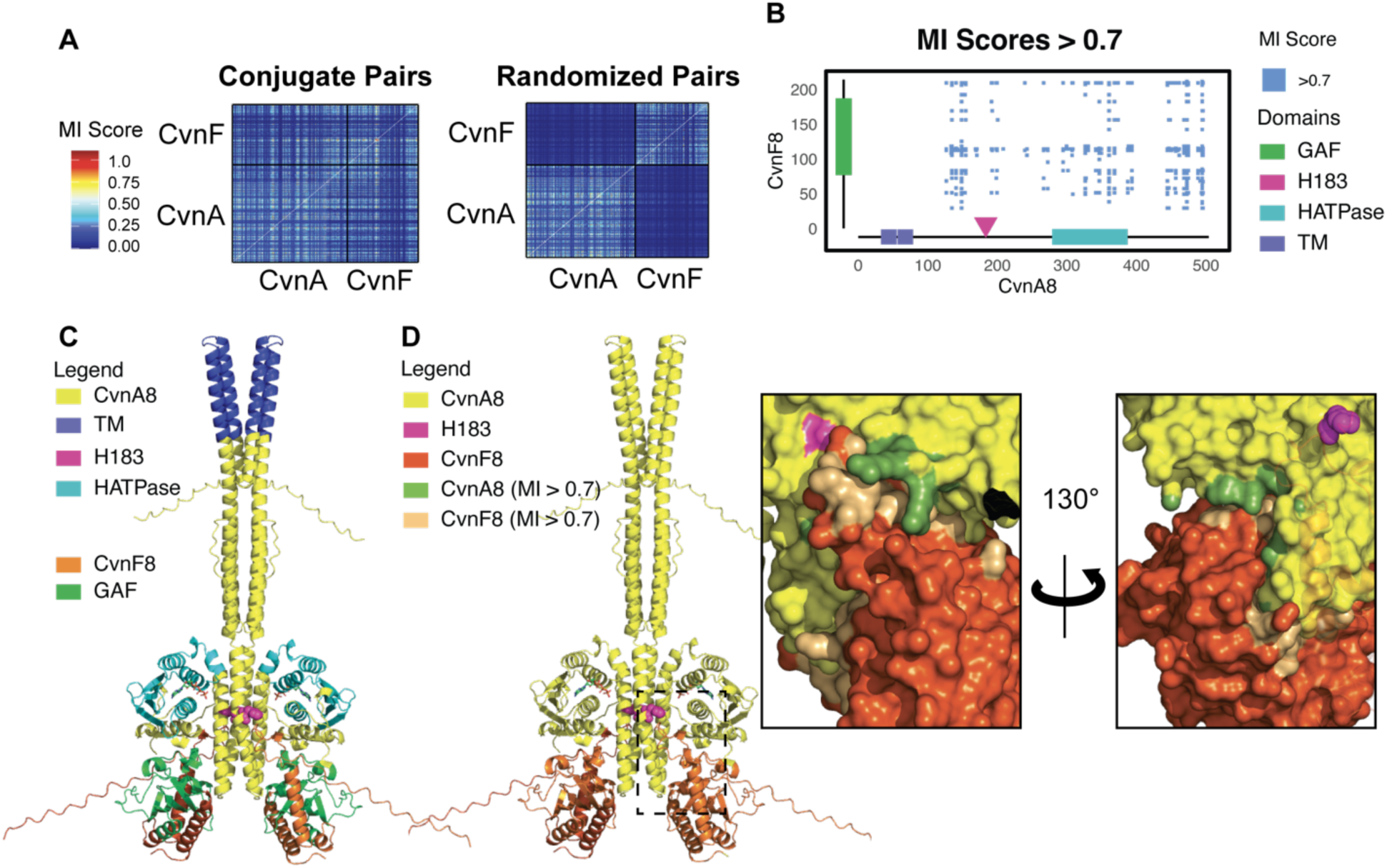
Mutual Information Analysis and Structural Context of Co-evolving Residues in CvnA and CvnF homologs. (A) Mutual Information (MI) scores for residue-residue coevolution between CvnA and CvnF proteins. The left heatmap displays MI scores for true conjugate pairs, and the right panel shows MI scores for randomized pairs. (B) MI scores above 0.7 mapped to CvnA8 and CvnF8 residues. PFAM annotations are represented as colored rectangles on the x and y axis. (C) AlphaFold structural models of CvnA8 and CvnF8 from *S. coelicolor*, with Pfam annotations. (D) AlphaFold3 model with MI scores mapped onto the protein structure (insets). Residues with high MI scores and close spatial proximity are highlighted, suggesting putative interaction interfaces. Additionally, a histidine residue (shown in magenta) is identified as a potential autophosphorylation site in CvnA8.

We next used AlphaFold3 to build a structural model of a CvnA8 dimer and two copies of CvnF8 from *Streptomyces coelicolor* (Fig. 3C). This structural modeling placed the two CvnF8 monomers at the base of the CvnA8 dimer. Each CvnF8 monomer has an interface directly in contact with the HATPase domains of CvnA8, suggesting potential protein-protein interactions.

We next mapped the between-protein coevolving residues from the MI analysis onto the AlphaFold3-predicted structural models to assess their spatial distribution (Fig. 3C, inset). Mapping high MI-scoring residues (MI > 0.7) in CvnA8 and CvnF8 showed that many coevolving sites are localized at the predicted interaction interface. Additionally, a highly conserved histidine residue (H183), which could serve as the site of CvnA8 autophosphorylation (highlighted in magenta in Fig. 3C), is located in proximity to the predicted coevolving interface.

Taken together, these findings indicate that CvnA and CvnF protein residues are evolutionarily constrained, and in the example of CvnA8 and CvnF8 from *S. coelicolor*, many of their coevolving residues are located at a predicted interaction interface near the likely site of CvnA autophosphorylation. These findings are broadly consistent with the idea that CvnF has coevolved as an accessory modulator of CvnA activity, with implications for control of CvnA autophosphorylation.

### CvnF8 enhances CvnA8 ATPase activity *in vitro*

Sensor histidine kinases hydrolyze ATP, yielding the phosphoryl group that is transferred to the receiving histidine in their DHp domains. This hydrolysis is carried out by globular HK ATPase domains, as noted in cyan in Fig. 3C. In typical sensor histidine kinases, ATP hydrolysis and autophosphorylation are triggered upon receiving a stimulus via their extracellular sensory domains. However, the modeling and analyses in the previous figures led us to hypothesize that CvnF8 might control CvnA8 ATPase activity and autophosphorylation through direct interactions.

To test this possibility, we measured ATP turnover using kinase luminescence assays (Fig. 4, Fig. S6). We first sought to measure ATP turnover by CvnA8 alone and in the presence of CvnF8, as a function of ATP substrate concentration. We also checked ATP turnover by a CvnA8 H183A mutant protein, which lacks the histidine residue that is a potential target of autophosphorylation. ATP hydrolysis by CvnA8 was measured at ATP concentrations ranging from 0 to 300 µM, and kinetic parameters were determined by nonlinear regression fitting to the Michaelis-Menten equation.

**Fig. 4.**
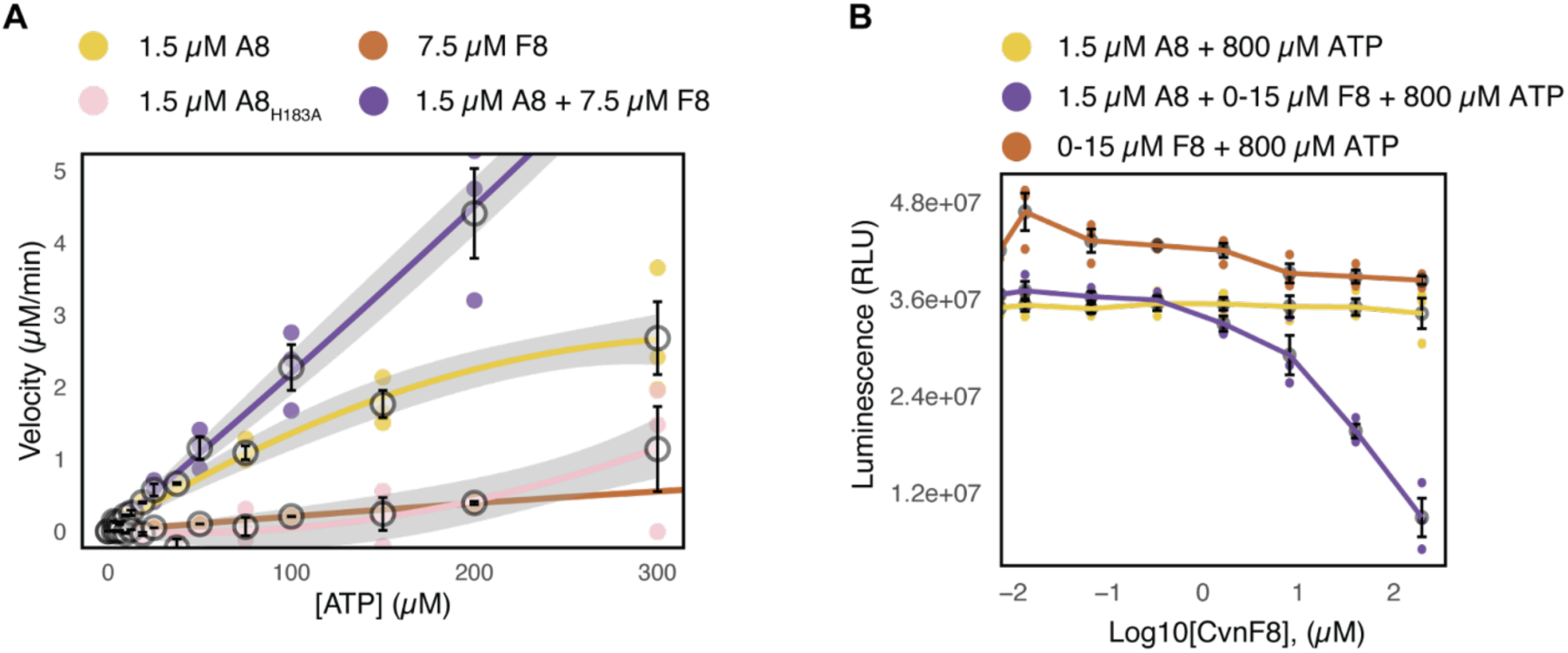
ATPase Activity of CvnA8 is stimulated by CvnF8 and depends on Histidine-183. (A) Michaelis-Menten kinetics of CvnA8 wild-type (A8) and the H183A mutant (H183A), showing a significant reduction in ATP turnover in the mutant. The disruption of activity in H183A supports the role of Histidine-183 as a key catalytic residue in CvnA8 function. Including CvnF8 together with CvnA8 leads to a large increase in ATPase activity (purple line) (B) Kinase-Glo assay measuring ATP consumption in reactions containing CvnA8 alone (A8), CvnA8 with increasing concentrations of CvnF8 (A8+F8), and CvnF8 alone (F8). Luminescence (RLU) is plotted against log-transformed CvnF8 concentration, revealing that higher levels of CvnF8 enhance ATP turnover, indicating a stimulatory effect on CvnA8 activity.

Wild-type CvnA8 displayed ATPase activity with a kcat of 0.044 s^-1^, indicating moderate ATP turnover (Table 1). The catalytic activity was drastically reduced in the H183A mutant, which prohibited the calculation of a nominal kcat (Fig. 4A). This finding confirmed that Histidine-183 is essential for normal ATP hydrolysis by CvnA8. In contrast, when CvnF8 was included with CvnA8, we found a kcat of 0.32 s^-1^, indicating a much higher turnover rate of ATP (Table 1). We attribute the turnover of ATP solely to CvnA8 in this system, as CvnF8 alone did not hydrolyze significant amounts of ATP. The maximum velocity (Vmax) of ATP hydrolysis was also significantly increased in the presence of CvnF8.

**Table 1.** Kinetic constants of ATP hydrolysis by CvnA8 alone and together with CvnF8. Michaelis-Menten constants were determined from nonlinear regression fits of initial velocity versus ATP concentration. Values represent the apparent Km, Vmax, kcat, and kcat/Km calculated from reactions using purified proteins.

| Protein | $K_m$ ($\mu\text{M}$) | $V_{max}$ ($\mu\text{M s}^{-1}$) | $k_{cat}$ ($\text{s}^{-1}$) | $k_{cat} / K_m$ ($\text{mM}^{-1} \text{s}^{-1}$) |
| --- | --- | --- | --- | --- |
| CvnA8 | $99.37 \pm 15$ | $0.032 \pm 0.004$ | $0.044 \pm 0.006$ | $4.43 \times 10^{-4} \pm 0.0001$ |
| CvnA8+CvnF8 | $1310 \pm 369$ | $0.32 \pm 0.06$ | $0.32 \pm 0.06$ | $2.45 \times 10^{-4} \pm 0.0001$ |

To further investigate the effect of CvnF8 on CvnA8, ATP turnover was measured in reactions containing 1.5 𝜇M CvnA8 alone, or with increasing concentrations of CvnF8 (Fig. 4B). In these experiments, the luminescence intensity is an indicator of ATP remaining after the reaction is stopped. Thus, lower luminescence indicates more ATP hydrolysis during the experiment. We found that the luminescence intensity (RLU) dramatically decreased when CvnF8 was included at a molar ratio of 2.5 CvnF8 to 1 CvnA8, or higher. This enhanced ATP turnover continued to increase if CvnF8 was included up to a ratio of 10:1, which was the highest ratio we tried. Taken together, these results indicate that CvnF8 can strongly potentiate the ATPase activity of CvnA8.

### CvnF8 enhances CvnA8 autophosphorylation *in vitro*

The biochemical experiments in the previous section prompted us to consider two hypotheses. Namely: A) that CvnA8 undergoes autophosphorylation at H183, and B) that CvnF8 can drive CvnA8 autophosphorylation. To test these hypotheses, we used Phos-Tag^TM^ gel shift assays to assess autophosphorylation over time, comparing wild-type CvnA8 and the H183A mutant, and CvnA8 in the presence of CvnF8 (Fig. 5). After electrophoresis, proteins were transferred to membranes for western blotting, and were detected with Strep antibody HRP. In these experiments, phosphorylation causes a mobility shift and is seen as additional protein bands that appear near the top of the blot images.

**Fig. 5.**
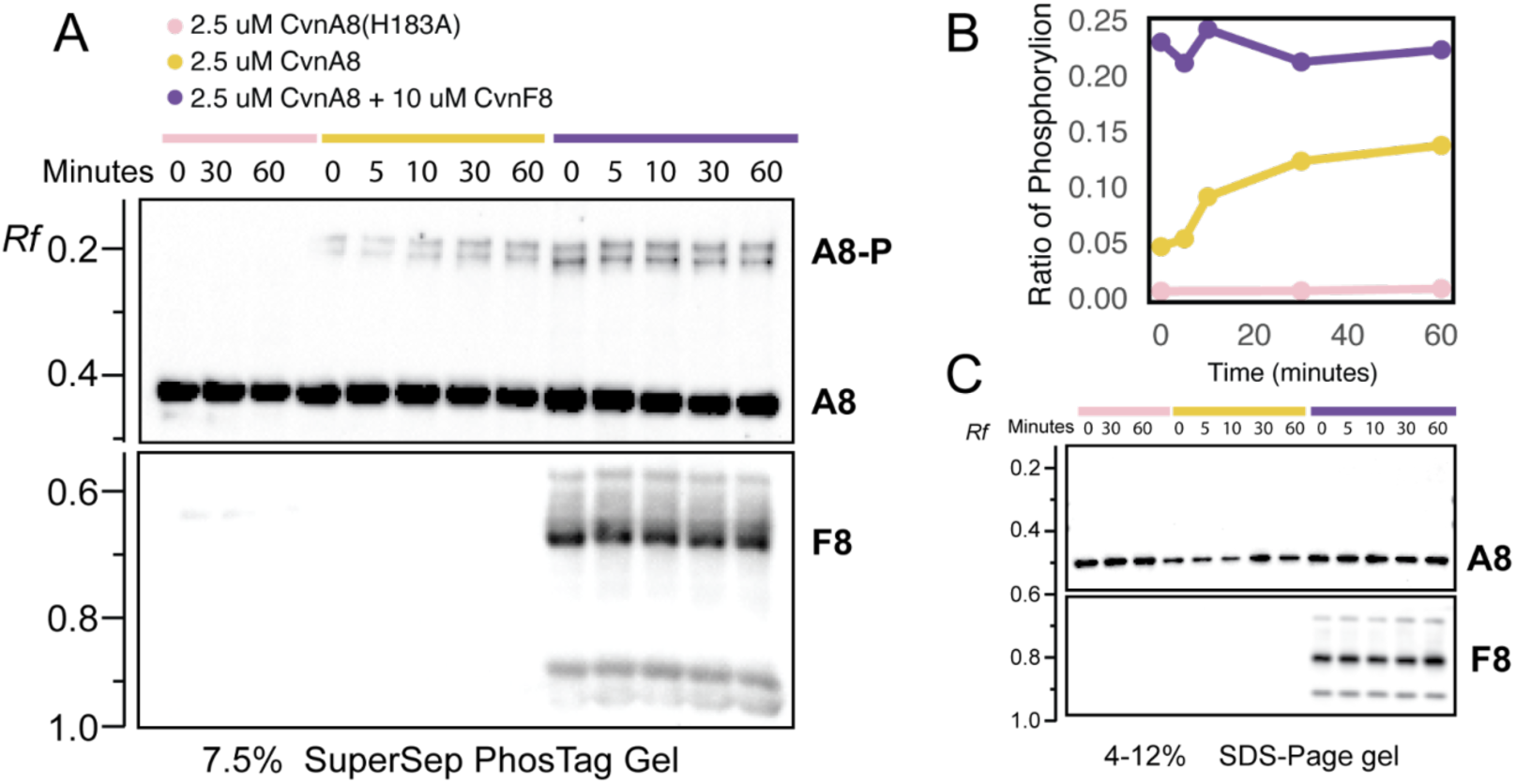
CvnA8 Autophosphorylation and Regulation by CvnF8. (A) Representative image of a western blot after Phos-Tag gel electrophoresis showing the mobility shift of CvnA8 due to autophosphorylation. All proteins bear a strep tag, and blots were probed with Strep antibody HRP. Conditions include the H183A mutant of CvnA8 (2.5 µM CvnA8:H183A) alone, CvnA8 alone (2.5 µM), and CvnA8 with CvnF8 (2.5 µM CvnA8 + 10 µM CvnF8). Robust autophosphorylation is seen when CvnF8 is included (lanes marked with purple bar). The H183A mutant lacks a detectable shift, confirming the essential role of Histidine-183 in autophosphorylation. (B) Quantification of phosphorylation over time. Phosphorylation levels were quantified as the ratio of phosphorylated to total protein and plotted as a function of time. The presence of CvnF8 increases the rate of CvnA8 autophosphorylation compared to CvnA8 alone. (C) Western blot of control SDS-PAGE electrophoresis with the same conditions as A. Bands in SDS-gel lack a detectable shift.

WT CvnA8 and the CnvA8 H183A mutant protein were incubated at discrete intervals including 0, 30, and 60 min for the H183A mutant and 0, 5, 10, 30, and 60 min for WT CvnA8 in buffer with excess ATP (see materials and methods). As seen in Fig. 5A, the WT CvnA8 protein showed a progressive mobility shift, indicating increasing autophosphorylation over time (middle lanes), such that by 60 minutes, autophosphorylation had plateaued at roughly 15% of the total CvnA8 protein (yellow plot in Fig. 5B). In contrast, the H183A mutant did not exhibit a shift, confirming that Histidine-183 is required for autophosphorylation.

To assess if CvnF8 altered CvnA autophosphorylation, CvnF8 was incubated with WT CvnA8 (at a ratio of 4:1 CvnF8 to CvnA8) in the same conditions as outlined above. The presence of CvnF8 rapidly increased the phosphorylation rate and extent of CvnA8, as seen in the rightmost lanes of Fig. 5A. Immediately after addition of CvnF8 (at 0 min), the percentage of phosphorylated CvnA8 was around 24%, compared to around 5% for CvnA8 alone (Fig. 5B).

With CvnF8 included, the amount of phosphorylation of CvnA8 remained stable at around 24% for the 60 min duration of the experiment. Together, these assays confirm that CvnA8 undergoes histidine autophosphorylation, Histidine-183 is essential for this activity, and that CvnF8 enhances CvnA8 autophosphorylation.

### Cvn8 system function and epistasis in double *cvnF8*/*cvnA8*/*cvnB8* mutants

We considered how the results above might be integrated with our current, limited understanding of how the Cvn8 system may function (Fig. 6A). This working model contains several unknowns; for example which Cvn8 components may be phosphorylated by CvnA8 and which downstream effector may serve as a DNA-binding protein for direct regulation of the *lan* gene cluster. However, this model remains useful for considering possible epistatic relationships between components of the Cvn8 system.

**Fig. 6.**
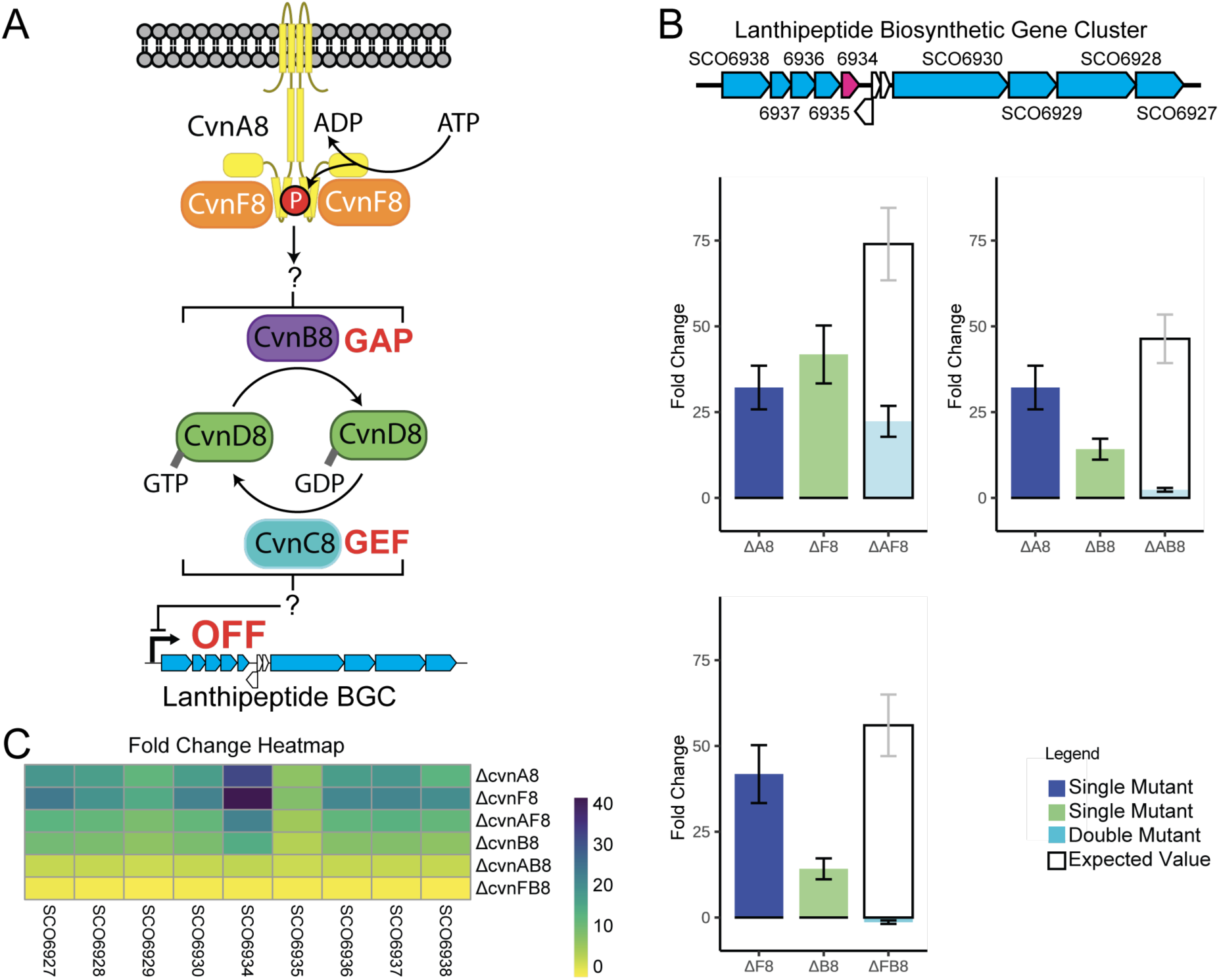
Working model of Cvn8 AGPS function and epistatic effects among Cvn8 components. (A) Model of the Cvn8 signaling system, in which CvnF8 drives ATPase activity/autophosphorylation of CvnA8. The target of phosphorylation by CvnA8 is unknown, but presumably influences the GTP-binding state of CvnD8. An unknown effector downstream of CvnD8 mediates repression of the *lan* BGC. (B) Bar graph showing average fold-change in expression of SCO6934 across single and double deletion mutants of Cvn8 genes (CvnA8, CvnB8, and CvnF8). The gene diagram at the top highlights the location of SCO6934 in magenta within the *lan* BGC. Genes colored blue were also checked via NanoString. Error bars represent standard deviation (n = 3). The rightmost hollow bar in each set represents the expected additive effect assuming no epistasis (i.e., independent gene function). (C) Heatmap of *lan* BGC gene expression changes across relevant mutants. Strong negative epistatic interactions were observed between all Cvn8 components tested.

To test for possible epistasis between proteins that we hypothesize are involved in the upper parts of the CvnA8 system cascade, we constructed several c*vn8* double mutant combinations and once again checked expression of genes in the neighboring lanthipeptide biosynthetic gene cluster using NanoString nCounter analysis, with similar growth conditions to those used in Fig. 2.

One prediction is that mutants lacking both CvnA8 and CvnF8 (a *ΔcvnAF8* strain) should show an expression pattern that indicates their epistatic interaction. Specifically, the derepression of the *lan* genes in a *ΔcvnAF8* strain should be nonadditive if they interact (10). When we checked the transcriptional output of a *ΔcvnAF8* strain, we found that genes of the *lan* gene cluster were indeed derepressed to a lower extent than expected if they functioned independently. For example, in the *ΔcvnAF8* strain, the *lan* gene SCO6934 was expressed 22.3-fold higher than the WT, falling far short of the expected value of 74-fold higher expression if CvnA8 and CvnF8 functioned independently (Fig 6B). This pattern of negative epistasis was true for expression across nine *lan* genes (heatmap in Fig. 6C and supplemental Fig. S3-S5). We note that, surprisingly, the nine *lan* genes we checked were all derepressed to a lower extent in the *ΔcvnAF8* strain compared to both the *ΔcvnA8* and *ΔcvnF8* strains (heatmap in Fig. 6C).

We also examined expression of the *lan* gene cluster in mutants lacking *cvnA8* or *cvnF8* in combination with *cvnB8*. We found that both *ΔcvnAB8* and *ΔcvnFB8* strains exhibited restored repression of the *lan* genes (Figs. 6B and 6C). This pattern of gene expression once again deviated from the expected additive effect if these components functioned independently. Taken together, these patterns of negative epistasis indicate interactions between CvnA8, CvnF8, and CvnB8.

## Discussion

Conservon systems are actinobacterial signal transduction systems composed of four core proteins, conventionally labeled CvnA-D. We propose that conservons should be understood as Actinobacterial G protein systems (AGPSs), alluding to the phylum-specific distribution and core G protein components of these systems. Some AGPSs have additional accessory proteins, including CvnF-types (GAF domain-containing proteins) and CvnE-types, which contain cytochrome P450 domains. Key questions remain about signal transduction within AGPSs, and what role might be played by accessory CvnF and CvnE proteins. Our findings indicate that accessory proteins likely serve as modulators of a subset of CvnA histidine kinases, placing them at the beginning of AGPS cascades.

### Expansion of AGPSs in a subset of Actinomycetota lineages

The broad but non-universal distribution of AGPSs across 485 phylogenetically diverse actinobacterial genomes suggests that the last common ancestor the Actinomycetota may have carried an AGPS, with subsequent lineage-specific losses shaping their current distribution. This hypothesis aligns with the analysis of Wuichet et. al. (2), which suggested that a RAS-like GTPase (i.e. the ancestor of CvnD/MglA homologs) was present in the last common ancestor of all bacteria. The authors suggest that these CvnD/MglA homologs were subsequently retained in only a few extant bacterial phyla, including the Actinomycetota, Myxococcota, and Bdellovibrionata.

Multiple actinobacterial clades with large genomes are enriched in AGPSs. This enrichment is in line with known correlations in genome size and the number of regulatory systems, wherein the larger the genome, the greater the proportion of genes dedicated to regulatory functions (11, 12). The abundance of AGPSs in these clades suggests that in these lineages, AGPSs have likely undergone duplication, diversification of function, and possible horizontal gene transfer.

### Modularity of CvnA histidine kinases and the distribution of CvnF and CvnE accessory proteins

CvnA histidine kinase homologs show sequence variability, leading to a variety of domain architectures as seen in Fig. 2A. The majority of CvnA homologs possess extracytoplasmic NIT-type sensory domains which are known to sense nitrate/nitrite (13, 14). Notably, examples of NIT domains exist that do not bind nitrate/nitrite and have no known ligands (15, 16). Given that at least six of the thirteen CvnA homologs in *S. coelicolor* have NIT domains, it seems likely that they might sense ligands other than nitrate/nitrite, although this remains to be tested.

A key observation from the analysis in Fig. 2 is that *cvnF* and *cvnE* accessory genes were reliably found with cognate CvnA homologs that lack NIT domains. Annotating CvnF sequences with PFAM indicated that these proteins contain a single GAF domain. GAF domains are known to bind a variety of ligands including cyclic nucleotides (17), amino acids (18), hemes (19), Fe-S clusters for oxygen sensing (20), and bilins for light sensing (21), often prior to exerting allosteric control of other protein domains.

GAF domains are also found in two component histidine kinases, though in these cases this domain is often positioned between the transmembrane domains and the DHp domain (22, 23). In the archetypal case of the potassium sensor KdpA, the GAF domain is thought to have a role in dimerization and/or signal transduction to the DHp domain, and any ligand binding by this domain remains undetermined (23). Within phosphodiesterases, GAF domains are known to bind cyclic nucleotides and subsequently modulate enzymatic activities of domains within the same protein (17, 24). We note that CvnF8 does not possess a conserved NKFDE motif, which is characteristic of cyclic nucleotide binding in the GAF domains of many phosphodiesterases (17, 24, 25), suggesting that its ligand is likely not a cyclic nucleotide.

We also note that CvnF accessory proteins contrast with the above examples in that they are stand-alone proteins whose structure is mostly composed of a GAF domain, and that lack other clearly catalytic domains (Fig 3C). We speculate that recruitment of CvnF accessory proteins by CvnA homologs may endow these sensors with two notable features: 1) it may enable them to sense cytosolic ligands, and 2) it may represent an evolutionary solution for sensing different ligand types, outside of the range of ligands afforded by the NIT domain.

### Modulation of CvnA8 activity by CvnF8

We used the Cvn8 system of *Streptomyces coelicolor* as a model to investigate the role of a CvnF-type accessory proteins in AGPS function. The structural model presented in Fig. 3 suggests that CvnF8 and CvnA8 may share an interface that is partially composed of coevolving residues. We hypothesize that CvnF8 might influence the position of the ATPase domains of CvnA8 such that autophosphorylation of histidine 183 is favorable. This is different from typical histidine kinases, which rely on relative shifts along the helical HAMP domains to bring the ATPase domains into proximity of the autophosphorylated histidine (26, 27). Further experiments will be needed to validate this model. To our knowledge, if confirmed, this interaction would represent a novel mechanism of histidine kinase activation.

The results presented in Figures 4 and 5 demonstrate that *in vitro*, CvnF8 enhances the ATPase/autophosphorylation activity of CvnA8. The catalytic constant of CvnA8 alone (kcat = 0.044 ± 0.006 s-1) falls within the range reported for other histidine kinases (0.001-0.083 s-1) (28–31). We measured the Km of CvnA8 for ATP to be 99.37±15 μM, which is also comparable to values observed across histidine kinases, which typically range from ∼10-200 μM (28). The presence of CvnF8 dramatically increased the maximum velocity (Vmax) of ATP hydrolysis (10-fold) and the kcat (7.3-fold). We interpret these enhanced levels of ATPase/autophosphorylation activity, stimulated by CvnF8, as corresponding to active signal transduction by CvnA8.

### A developing picture of Cvn8 AGPS function

Our *in vitro* experiments leave open the question of whether CvnF8 exerts its activity on CvnA8 constitutively, or requires a stimulus *in vivo*. If CvnF8 potentiates CvnA8 activity in a constitutive manner, it is possible that binding of a ligand by the GAF domain of CvnF8 could lead to a cessation of CvnA8 autophosphorylation. This would interrupt signal flow through the rest of the Cvn8 AGPS. A second possibility is that *in vivo*, CvnF8 requires ligand binding to exert its activity on CvnA8. In this case, strong autophosphorylation of CvnA8 would only happen in the presence of the ligand recognized by the CvnF8 GAF domain. Differentiating between these possibilities will be challenging until the ligand for CvnF8 is determined.

As shown in Fig. 6A, the target of phosphorylation by CvnA8 remains unknown, but could be any of the remaining Cvn8 system proteins. Among these, we suggest that CvnB8 or CvnC8 are likely targets since altering either of their activities would provide a means of shifting the state of CvnD8 toward the GDP or GTP-bound form. One possibility is that the GTP-bound form of CvnD8 mediates the downstream repression of the *lan* biosynthetic gene cluster through interacting with an unknown DNA-binding protein. If this is true, it may explain why *ΔcvnAB8* and *ΔcvnFB8* strains exhibited restored repression of the *lan* biosynthetic gene cluster (Fig. 6B). One interpretation of these results is that in these double mutants, CvnC8 maintains CvnD8 in a near-completely GTP-bound state, restoring the aberrant derepression seen in single *ΔcvnA8*, *ΔcvnF8*, and *ΔcvnB8* strains. We emphasize that this model is a work-in-progress that may serve as a roadmap for future testing of its individual features and component interactions.

### Concluding remarks

The Cvn8 system was originally identified as a regulator of specialized metabolism in *S. coelicolor* during interactions with other actinomycetes (6). Our prior work implicated *cnvF8* in the function of the Cvn8 AGPS, but how CvnF8, and other accessory proteins, might interact with other AGPS components remained unclear. The present work demonstrates that CvnF8 can interact with, and modulate the autophosphorylation of, the CvnA8 histidine kinase.

Additionally, we show that a mutant lacking both CvnF8 and CvnA8 displayed patterns of gene expression that imply functional epistasis, suggesting that their function within the Cvn8 cascade is interdependent. Together, these results provide a function for CvnF8 accessory proteins, which are widespread in AGPSs found across diverse members of the Actinomycetota.

## Materials and methods

### Species Tree Construction

A species tree was constructed using 485 genomes selected from ActDES (Actinobacterial Database for Evolutionary Studies) (9), a curated, non-redundant database of *Actinomycetota* genomes. These genomes are publicly available and were downloaded from the Integrated Microbial Genomes (IMG) database (32). Phylogenetic inference was based on a concatenated alignment of 15 conserved ribosomal proteins: L2, L3, L4, L5, L6, L14, L15, L16, L18, L22, S3, S8, S10, S17, and S19. Multiple sequence alignment was performed using MAFFT v7.520 (33), employing the FFT-NS-i method. Poorly aligned regions were removed using ClipKit v1.3.0 (34) with the default smart-gap setting to retain informative sites while filtering out low-confidence regions. The resulting concatenated alignment was used to construct a maximum-likelihood phylogeny with IQ-TREE2 v1.6.12 (35), using the LG substitution model. Ultrafast bootstrap analysis was performed with 1,000 replicates. The final tree was visualized using ggtree v3.2.1 (36) within the ggplot2 (37) framework in R.

### Orthogroup Identification

The same set of 485 genomes from ActDES was used to identify protein families corresponding to conservon-associated orthogroups. Orthologous protein families were identified using OrthoFinder v2.5.5 (38), grouping proteins into orthogroups based on sequence similarity and phylogenetic inference. AGPS-associated orthogroups (*cvnA, cvnB, cvnC, cvnD, cvnE, cvnF*) were extracted, and their syntenic conservation across genomes was assessed by examining their genomic neighborhood. The presence or absence of each orthogroup in synteny for each genome was encoded as a binary matrix and visualized as a heatmap alongside the species tree using gheatmap from the ggtree package in R.

### Domain Annotation and Synteny Analysis

Protein sequences from the CvnA orthogroup were annotated using InterPro against the Pfam, and TMHMM databases to identify Transmembrane (TM), Nitrate Sensing (NIT), HAMP, and HATPase domains (39). The CvnA gene tree, generated by OrthoFinder, was visualized with ggtree, and a heatmap of domain presence was overlaid using gheatmap in R. To assess syntenic conservation, the presence of CvnE and CvnF was determined based on their genomic proximity to CvnA, requiring that CvnE or CvnF be within four genes of CvnA in the same genomic neighborhood. A binary matrix encoding the presence or absence of these syntenic genes was generated and displayed as an additional heatmap track aligned to the CvnA gene tree.

### Mutual Information Analysis

Multiple sequence alignments were generated for CvnA and CvnF orthogroups using MAFFT v7.520 with the FFT-NS-i method. Poorly aligned regions were removed using ClipKit v1.3.0 with the default smart-gap setting. Mutual Information (MI) scores were calculated using the mutualinfobio R library (https://github.com/TraxlerLab/mutualinfobio), which computes mutual information scores between aligned residues. MI scores were computed for CvnA-CvnF conjugate pairs and for randomized protein pairings as a control. MI scores were visualized as heatmaps using ggplot2 in R.

### Structural Mapping with AlphaFold

Protein structure predictions for CvnA8 dimers and CvnF8 monomers were generated using **AlphaFold3**(40) with default parameters. MI scores were mapped onto the predicted structures to visualize coevolving residues. PyMOL v3.1.3.1 was used to generate structural representations.

### Bacterial Strains and Plasmid Construction

Bacterial strains, and primers used in this study are listed in Tables S1-S2 respectively. All PCR reactions were performed using NEB Q5 High-Fidelity DNA Polymerase (NEB #M0491S) according to the manufacturer’s protocols and online tools. Plasmid construction was carried out using Gibson Assembly (NEB #E2611L) with two- or three-fragment assembly strategies.

The pET21-Strep-CvnA8 plasmid was constructed by assembling three fragments: (1) the cvnA8 coding sequence (nucleotides 55-518) was amplified from the St1G8 cosmid using primers LCRJ21-LCRJ04, introducing a 3x Strep tag at position 55, past the transmembrane domains, to facilitate protein purification; (2) the pET21 backbone was amplified using primer pairs LC2101-BB82 and BB98-LC2102, where BB82 and BB98 fully overlapped, and LC2101 and LC2102 had 17 bp overlaps with the LCRJ21-LCRJ04 fragment.

Similarly, the pET21-Strep-CvnF8 plasmid was constructed using three-fragment Gibson Assembly. The cvnF8 (SCO6939) gene was amplified from the St1G8 cosmid using primers LC2110-BB319, appending a 3x Strep tag to the N-terminus of the sequence. The pET21 backbone was amplified using LC211-BB82 and BB98-LC2102, following the same overlap-based assembly strategy.

Following PCR amplification, DNA fragments were separated on agarose gels, extracted using the Monarch DNA Gel Extraction Kit (NEB #T1020), and assembled via Gibson Assembly. The assembled plasmids were transformed into *E. coli* XL1-Blue cells via heat shock and plated on LB-agar plates with ampicillin (100 µg/mL) at 37°C overnight. Ampicillin-resistant colonies were screened, and plasmids were sequenced using Plasmidsaurus (Oxford Nanopore Technology) with custom analysis and annotation. Confirmed plasmids were transformed into E. coli BL21 for protein expression and purification.

*Streptomyces coelicolor* gene knockouts were generated using the Redirect PCR-targeting system.(41) First, the *cvnA8* gene was replaced in St1G8 with an Apramycin resistance cassette, followed by cassette removal using the pCP20 plasmid. Additional knockouts of *cvnB8* or *cvnF8* were performed using the same strategy. Streptomyces coelicolor M145 strains were cultured on ISP2 or MS agar plates following standard protocols.

### Expression and Purification of CvnA8 and CvnF8

Expression and purification of CvnA8 and CvnF8 were performed as previously described (Skerker et al., 2008). *E. coli* BL21 (DE3) cells harboring pET21-Strep-CvnA8 or pET21-Strep-CvnF8 were grown in Terrific Broth Autoinduction Medium (Formedium #AIMTB0210) at 30°C until reaching an OD₆₀₀ of 0.5, then moved to 19°C for overnight induction.

Following induction, cells were harvested by centrifugation, cells were resuspended in lysis buffer (25 mM Tris, 300 mM NaCl, pH 8) supplemented with a protease inhibitor cocktail (Complete, Roche), and stored at −20°C. Frozen cells were thawed on ice and lysed by two passages through a French Press using a large pressure cell at 30,000 psi. The lysate was centrifuged at 20,000 rpm for 30 minutes at 4°C, and the soluble fraction was filtered through 0.45 µm filters before purification.

Proteins were purified using affinity chromatography on a 5 mL Strep-Tactin XT column (Cytiva # 29401322) pre-equilibrated with buffer A (25 mM Tris, 300 mM NaCl, pH 8). After washing with buffer A, proteins were eluted with buffer A containing 50 mM biotin. Fractions were further purified using size-exclusion chromatography (SEC) on a Superdex 75 Increase 10/300 GL column (Cytiva) equilibrated in Storage Buffer (25 mM Tris, 50 mM NaCl, 10% glycerol, pH8).

Purity was assessed by SDS-PAGE with Coomassie staining, and protein concentration was determined using a Pierce Detergent-Compatible Bradford Assay Kit (ThermoFisher) following the manufacturer’s instructions. Proteins were concentrated using Amicon Ultra-3K centrifugal filters (Millipore, 3 kDa MWCO) prior to storage.

### Michaelis-Menten Kinetics Assay

ATP turnover kinetics was measured in triplicate using the Kinase-Glo Plus Luminescence Assay (Promega #V3771) in 384-well white plates (Greiner bio-one #5678-1075) with a final reaction volume of 10 µL. Reactions contained 25 mM Tris (pH 8.0), 150 mM NaCl, and 100 mM MgCl₂, with a fixed concentration of protein. (1.5 µM CvnA8, 1.5 µM CvnA8:H183A, 7.5 µM CvnA8, 1.5 µM CvnA8 with 7.5 µM CvnF8). For all the alone conditions, ATP was serially diluted 1:2 from an initial concentration of 300 µM to 0 µM to generate a substrate saturation curve. For the CvnA8 with CvnF8 condition, ATP was initially serially diluted 1:2 from an initial concentration of 800 µM to 0 µM, but this ATP range did not reach substrate saturation. To calculate kinetic constants of the CvnA8 and CvnF8 complex a much higher concentration range (2.5 mM to 0) was necessary and then diluted 1:300 to measure the reaction. All reactions were incubated at 37°C for 30 minutes, followed by the addition of 10 µL Kinase-Glo reagent. Luminescence was measured after 10 minutes at room temperature using an EnVision 2104 Plate Reader (PerkinElmer). Data were analyzed in R, and ATP turnover rates were fitted to the Michaelis-Menten equation using nonlinear regression (nls function) to determine the kinetic constants Km, Vmax, kcat, and kcat/Km.

### Kinase-Glo Assay for CvnA8 and CvnF8 Interactions

ATP consumption was measured using the Kinase-Glo Luminescence Assay (Promega #V3771) in 384-well white plates (Greiner bio-one #5678-1075) with a final reaction volume of 10 µL. Each reaction contained 25 mM Tris (pH 8.0), 150 mM NaCl, and 100 mM MgCl₂, 800 µM of ATP, with 1.5 µM CvnA8 alone or in the presence of CvnF8 at final concentrations ranging from 0 to 15 µM. Reactions were incubated at 30°C for 30 minutes, followed by the addition of 10 µL Kinase-Glo reagent. After 10 minutes of incubation at room temperature, luminescence was measured using an EnVision 2104 Plate Reader (PerkinElmer). Luminescence values (Relative Light Units, RLU) were normalized to a no-enzyme control, and data was plotted in R to assess ATP consumption based on relative luminescence intensity across increasing concentration of CvnF8.

### Phos-Tag Gel Electrophoresis

Protein phosphorylation was analyzed using SuperSep Phos-Tag gels (Fujifilm #193-17981, 50 µM Phos-Tag, 7.5% acrylamide, 17-well format). Autophosphorylation reactions were performed by incubating 2.5 µM of protein with 800 µM ATP at discrete time points (0-60 minutes), followed by quenching reactions with Laemmli sample buffer without heat denaturation before loading. Electrophoresis was performed at 20 mA at 4°C until the Bromophenol Blue (BPB) dye front reached the bottom of the gel. Following electrophoresis, gels were soaked in Transfer buffer (192 mM glycine, 25 mM Trisbase, 20% methanol, pH 8.0) containing 5 mM EDTA for 10 minutes to remove excess Zn²⁺, ensuring efficient protein transfer for western blot analysis (see below).

### Western Blot Analysis

Proteins were transferred to PVDF membranes using the Trans-Blot Turbo system (Bio-Rad) according to the manufacturer’s instructions. Membranes were blocked in 5% non-fat milk in PBST (Phosphate buffered saline pH 7.2 with 0.5% Tween-20) for 1 hour at room temperature, followed by incubation with Strep antibody HRP (Cube Biotech) overnight at 4°C. After washing in PBST, membranes were developed using chemiluminescence (ECL) and imaged on a Bio-Rad ChemiDoc system. Phosphorylation levels were quantified by measuring the ratio of phosphorylated to total protein and analyzed using ImageJ.

### Transcriptomic analysis of Epistasis effects on cvn operon

RNA samples were prepared by growing patches of *S. coelicolor* wild-type (WT) and double-knockout mutant strains on agar rich media as previously described (42). Briefly, 0.5 µL of spore stock solutions were spotted onto ISP2 agar plates and incubated at 30°C for 3 days. Whole cells collected with a cell scraper from ISP2 plates and immediately ground in liquid nitrogen using a mortar and pestle. After the liquid nitrogen had evaporated, 150 µl of buffer RLT (Qiagen) was added to the mortar and the biomass was further pulverized to yield a crude cell lysate. 0.5 µl of this lysate was subjected to probe hybridization and processing with the Nanostring nCounter Spring Station according to the manufacturer’s instructions. Raw code counts were analyzed according to manufacturer’s guidelines; briefly, total transcript counts were normalized using internal controls with background subtraction. Transcript counts for three genes (*hrdB* , *folB* , and *hemL*) were used for geometric mean normalization to correct for differences in total mRNA concentration. All data were collected from three biological replicates, and fold changes were calculated for single mutants vs. wild-type and double mutants vs. wild-type. In Figure 6, the right hollow column represents the expected sum of average transcript fold-change under the assumption of independent gene function, with error bars calculated by propagating the standard error from single mutant fold changes.

## Acknowledgements

We thank Dr. Bailey Bonet for providing cvn8 mutant strains and for sharing the pET21 expression plasmid. We also thank Dr. Marie Elliot of McMaster University for the *S. coelicolor* cosmid used to build *S. coelicolor* mutants in this study. We acknowledge Drs. Kathleen Ryan, Ceci Martinez-Gomez and Adam Deutschbauer for discussion of experiments. We are grateful to Dr. Arash Komeili for access to an ÄKTA Pure chromatography system. We thank Emanuela Zacco of the HDFCCC Laboratory for Cell Analysis Shared Resource Facility at UCSF for her assistance with processing samples for NanoString data collection; this facility is supported by NIH grant P30CA082103. This research used the Savio computational cluster resource provided by the Berkeley Research Computing program at the University of California, Berkeley (supported by the UC Berkeley Chancellor, Vice Chancellor for Research, and Chief Information Officer). Funding for this work was provided by NIH R35GM128849 awarded to Prof. M. F. T. L. C. acknowledges support from the National Science Foundation Graduate Research Fellowship Program.

## Author Contributions

L.C., and M.F.T. designed experiments. L.M.C., and K.M. built experimental strains. K.D. technical assistance with protein purification. L.M.C. conducted experiments. L.M.C., and M.F.T. analyzed the data. L.M.C., and M.F.T. wrote the paper.

